# Divalent cations can control a switch-like behavior in heterotypic and homotypic RNA coacervates

**DOI:** 10.1101/453019

**Authors:** Paulo L. Onuchic, Anthony N. Milin, Ibraheem Alshareedah, Ashok A. Deniz, Priya R. Banerjee

## Abstract

Liquid-liquid phase separation (LLPS) of RNA-protein complexes plays a major role in the cellular function of membraneless organelles (MLOs). MLOs are sensitive to changes in cellular conditions, such as fluctuations in cytoplasmic ion concentrations. To investigate the effect of these changes on MLOs, we studied the influence of divalent cations on the physical and chemical properties of RNA coacervates. Using a model arginine-rich peptide-RNA system, we predicted and observed that variations in signaling cations exert interaction-dependent effects on RNA LLPS. Changing the ionic environment has opposing effects on the propensity for heterotypic peptide-RNA and homotypic RNA LLPS, which results in a switch between coacervate types. Furthermore, divalent ion variations continuously tune the microenvironments and fluid properties of heterotypic and homotypic droplets. Our results may provide a generic mechanism for modulating the biochemical environment of RNA coacervates in a cellular context.

## Introduction

Phase separation aids in regulating essential functions of cells and organisms. Within the cellular context, condensation-driven phase separation of biomolecules selectively compartmentalizes essential proteins, nucleic acids, and biochemical processes^1^. Liquid-liquid phase transitions can drive dynamic intracellular compartmentalization to form membrane-less organelles (MLOs) such as the nucleolus and stress granules^2–5^. Unlike their membrane-bound counterparts such as the nucleus, the unique physicochemical features of MLOs can enable liquid-like behavior such as facile formation, fusion, and dissolution^2^. These dynamic qualities provide a means for cells to sense and respond rapidly to changing environments, such as the cytoplasm during stress^2^. In this context, stimulus-dependent liquid-liquid phase separation (LLPS) of biopolymers has recently emerged with numerous implications in biology^6,7^.

One of the driving forces of MLO formation is the weak multivalent interactions among proteins and nucleic acids^8^. Quantitative understanding of the underlying interaction network and elucidation of molecular driving forces that alter them are therefore key topics of research. The interchain associations of proteins and nucleic acids are largely determined by their primary sequences^8^. Intrinsically-disordered proteins (IDPs) with low-complexity repetitive sequences have been identified as drivers of MLO biogenesis, with charged sequences being one of the most common motifs found in intracellular MLOs^9^. Phase separation of charged low-complexity motifs are largely driven by electrostatic interactions^8,10,11^. This phenomenon, commonly known as complex coacervation, can be recapitulated in an *in vitro* model consisting of an arginine-rich peptide and RNA mixtures^12,13^. Our recent work demonstrated that a peptide-RNA mixture can display reentrant phase behavior, where droplets can form and dissolve due to monotonic variation of ionic peptide-RNA ratios alone^12^. This type of behavior can be modeled simply using early polymer chemistry theories^14,15^, with electrostatic interactions being the only effective interaction parameter. In addition to complex coacervation of RNA and IDPs, recent reports have observed liquid-liquid phase transitions *via* homotypic interactions of RNA chains in the absence of protein^16,17^.

The global phase behavior of ternary IDP-RNA systems is determined by a complex interplay between homotypic (IDP-IDP and RNA-RNA) and heterotypic (IDP-RNA) interactions. Hence, IDP-RNA mixtures can be sensitive to small changes in solution conditions such as temperature, ionic environment, pH, and IDP-RNA stoichiometry^8^. These alterations in physicochemical conditions can prompt dynamic responses in phase behavior by influencing the relative strengths and interplay of homotypic and heterotypic interactions. We reason that this interplay raises the possibility of an alternative stimulus-dependent LLPS modulation, *via* tuning of the relative magnitudes of competing interactions by solvent components. While complex coacervates are destabilized by cations due to charge shielding, RNA structures and interactions are stabilized by cations^18,19^, which should promote RNA-RNA homotypic phase separation.

The cytoplasmic concentrations of divalent ions are among the most regulated conditions in the cell^20–25^. Under rapidly changing cellular conditions, such as during stress, the flux of signaling cations coincides with MLO formation and dissolution dynamics^21,22,24,25^. Several recent studies exemplify the critical nature of divalent ions and their relationship to phase separation. One study shows that the phase separation of a stress granule-related IDP, TIA-1, can be regulated by Zn^2+^ ^26^. Another study revealed heat-induced phase separation of single stranded DNA in the presence of divalent cations^27^. Given the importance of divalent cations in MLO dynamics, we set out to determine how fluctuations in divalent ion concentrations could control the phase behavior of proteins and RNA. Our hypothesis is founded on the potential of divalent cations, such as Mg^2+^ and Ca^2+^, to have a destabilizing effect on heterotypic IDP-RNA interactions, while stabilizing homotypic RNA interactions. We anticipate that these opposing effects could give rise to a switch-like behavior with the capacity to regulate between complex coacervates (heterotypic) and RNA coacervates (homotypic) by modulating cation concentration in solution. Additionally, the same underlying opposing effects on molecular interactions could also result in continuous tuning of droplet physical properties and composition.

### A divalent cation has opposing effects on heterotypic (peptide-RNA) and homotypic (RNA-RNA) droplets

We previously studied important biophysical aspects (reentrant phenomena and non-equilibrium sub-compartmentalization) of LLPS using an *in vitro* model IDP-RNA system, consisting of an arginine-rich peptide {RP_3_ (RRxxxRRxxxRRxxx)} and a long homopolymeric single-stranded RNA {polyU (800-1000 kD)}^12^. This peptide-RNA system was chosen because it has been shown to be a valuable model for understanding the electrostatically-mediated phase behavior of intracellular ribonucleoprotein (RNP) granules^13^, which often contain IDPs featuring arginine-rich motifs^9^. Our previous studies showed that monovalent (Na^+^) cations reduce the propensity for phase separation in this system, which is consistent with the notion that heterotypic electrostatic interactions drive LLPS in such RNP systems^12^.

Here, we investigated the effects of divalent cations on the phase behavior of this system. Turbidity measurements (Fig 1a), along with laser scanning confocal fluorescence microscopy at different salt concentrations (Fig 1b) confirmed that increased concentration of divalent (Mg^2+^) cations reduced LLPS. This is expected due to a weakening of heterotypic electrostatic interactions between RP3 and RNA that drive complex coacervation of these oppositely charged biopolymers. This effect is similar to that of monovalent (Na^+^) cations (Fig S2), though divalent ions are more effective.

**Figure 1:**
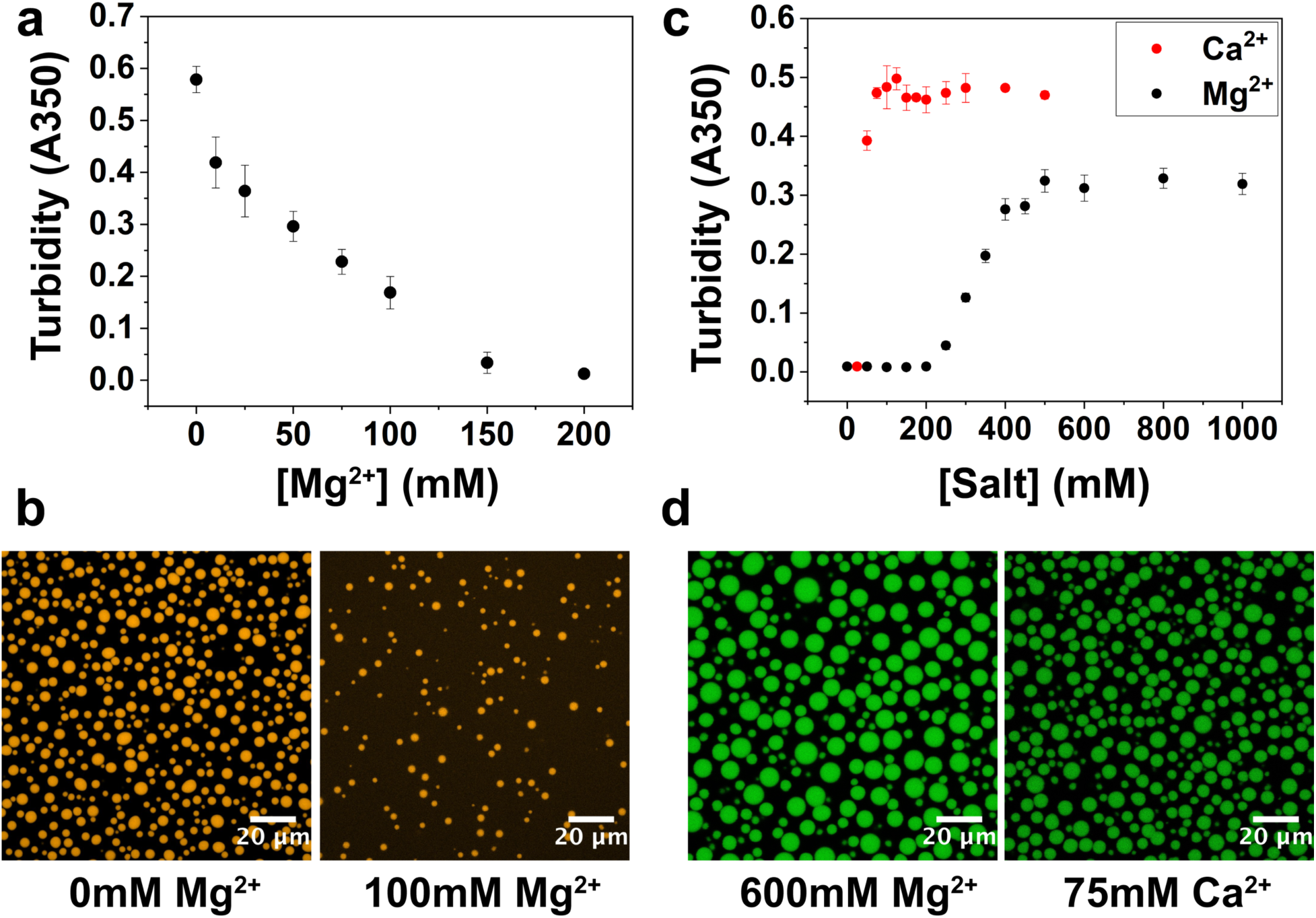
Increase in divalent cation concentration leads to dissolution of heterotypic RP3-polyU droplets and formation of homotypic polyU droplets. **a.** Solution turbidity measurements of RP3-polyU droplets as a function of [MgCl_2_]. ([RP3] = 500µM, 0.6x polyU wt/wt) **b.** Confocal fluorescence microscopy images of RP3-polyU droplets under different divalent salt conditions. ([RP3] = 500µM, 0.6x polyU wt/wt, 0.5µM RP3-AF594) **c**. Solution turbidity measurements of polyU coacervates as a function of [MgCl_2_] and [CaCl_2_] (1mg/mL polyU) **d.** Confocal fluorescence microscopy image of polyU coacervates (1.5mg/mL polyU, 600mM MgCl_2_, 0.5µM FAM-UGAAGGAC) (1.5mg/mL polyU, 75mM CaCl_2_, 0.5µM FAM-UGAAGGAC)

We then tested the influence of divalent cations on homotypic phase separation of RNA. It is well known that the structure and assembly of both RNA and DNA are heavily influenced by the presence of divalent cations^18,28^. This has been especially well-studied for Mg^2+^, which assists in the stabilization of RNA 3D structure, and has long been used in RNA folding and structural studies^18,28,29^. Therefore, in the case of polyU RNA, we anticipated that increased Mg^2+^ concentration could shield the electrostatic repulsion between the phosphate backbone of RNA and facilitate non-canonical U-U base pairing interactions^30–32^, thus stabilizing droplet formation. Indeed, it has been shown that spermine and spermidine, which are small organic cations with 4+ and 3+ charges, respectively, can lead to polyU coacervation^33^. A recent report also demostrated that varying amounts of monovalent cations (Na^+^) and a molecular crowder (PEG) can lead to phase separation of polyU^17^. In our system, we tested if variation of divalent cation concentration was sufficient to induce polyU coacervation in the absence of PEG. Using turbidity measurements, we investigated LLPS of polyU upon increasing Mg^2+^ and Ca^2+^ (Fig 1c), along with Sr^2+^ and Zn^2+^ (Fig S3). Solution turbidity data in conjunction with confocal fluorescence microscopy (Fig 1c, 1d, Fig S3, S4) showed that polyU forms droplets in the presence of Mg^2+^ (≥ 300mM), Ca^2+^ (≥ 50mM) and Sr^2+^ (≥ 75mM) in a concentration dependent manner, with Ca^2+^ and Sr^2+^ showing a lower concentration threshold required for phase separation. One potential explanation for this discrepancy between the phase separation thresholds of Mg^2+^ and Ca^2+^/Sr^2+^ is the difference in ion charge density, as it is known to have a profound influence on RNA stability^34^. Ca^2+^ and Sr^2+^ are more effective at screening the negative charge of the RNA phosphate backbone and have weaker nonspecific interactions with RNA than Mg^2+^. This difference is also consistent with previous data that showed a lower temperature threshold for ssDNA phase separation with Ca^2+^ than with Mg^2+^, but there is a difference in that phase separation was not observed in ssDNA in the presence of Sr^2+^ ^27^. In our work, the condensates displayed liquid-like characteristics such as fusion, circular appearance, and recovery of fluorescence after photobleaching (Fig. 1, 4, S3, S4). Zn^2+^ displayed different behavior in that it caused aggregation at high concentration (500mM), consistent with previous data^27^, but showed liquid-liquid phase separation at lower concentrations (≥ 75mM – 150mM) (Fig S4). The homotypic polyU droplets in the presence of 150mM Zn^2+^ displayed all of the same liquid-like characteristics as the droplets with the other ions, meaning that there is a concentration window with Zn^2+^ where the droplets are liquid-like prior to the observed aggregation (Fig S4). Addition of PEG substantially lowers the phase separation threshold concentration for polyU and Mg^2+^ from 300mM Mg^2+^ (0% PEG) to 75mM Mg^2+^ (5% PEG) to 50mM Mg^2+^ (10% PEG) (Fig S3). This is consistent with the known effect of PEG in nucleic acid cluster formation^35–37^. We also observed polyU phase separation at ≥ 400mM Na^+^ in the presence of 10% PEG (Fig S3). This suggested that the multivalency of the cation is not necessary to induce phase separation, as Na^+^ is a monovalent cation, but it does dramatically alter the phase separation threshold of polyU. The fact that high concentrations of Na^+^ and 10% PEG were required for the phase separation of polyU is consistent with monovalent ions having considerably smaller charge screening propensity and influence on RNA than divalent ions^18,19^.

### Divalent salt triggers a switch-like phase behavior of a peptide-RNA system

The above results clearly demonstrate the opposing effects of divalent cations on homotypic and heterotypic phase separation in a peptide-RNA system. If the phase boundaries (Fig 1a, 1c) remain consistent in the combined RP3-polyU system, Mg^2+^ should be able to act as a stimulus to create a switch-like phase behavior. In other words, starting with RP3-polyU droplets, a titration of Mg^2+^ would first lead to dissolution of these heterotypic droplets and subsequent formation of homotypic RNA droplets. Sequential turbidity measurements of RP3-polyU mixtures over increasing [Mg^2+^] corroborated this prediction, displaying a window of miscibility between the two-phase regions of the phase diagram (Fig 2a). To more directly visualize switching between different regions of the phase diagram, we used confocal fluorescence microscopy with differentially labeled RP3 (orange) and polyU (green) (Fig 2c). Starting with heterotypic RP3-polyU coacervates at low salt (10mM Tris, 0mM Mg^2+^), we added 150mM Mg^2+^ into the solution, which resulted in the dissolution of the droplets. Subsequent addition of Mg^2+^ to a final concentration of 500mM produced new homotypic polyU droplets (Fig 2c). Given the orthogonal effects of Mg^2+^ on our system, we can conclude that these are two distinct types of polyU RNA droplets. These two types of droplets will henceforth be referred to as heterotypic RP3-polyU droplets and homotypic polyU droplets based on these two types of droplet regimes described above. We also characterize the properties of the two droplet types in later sections below.

**Figure 2:**
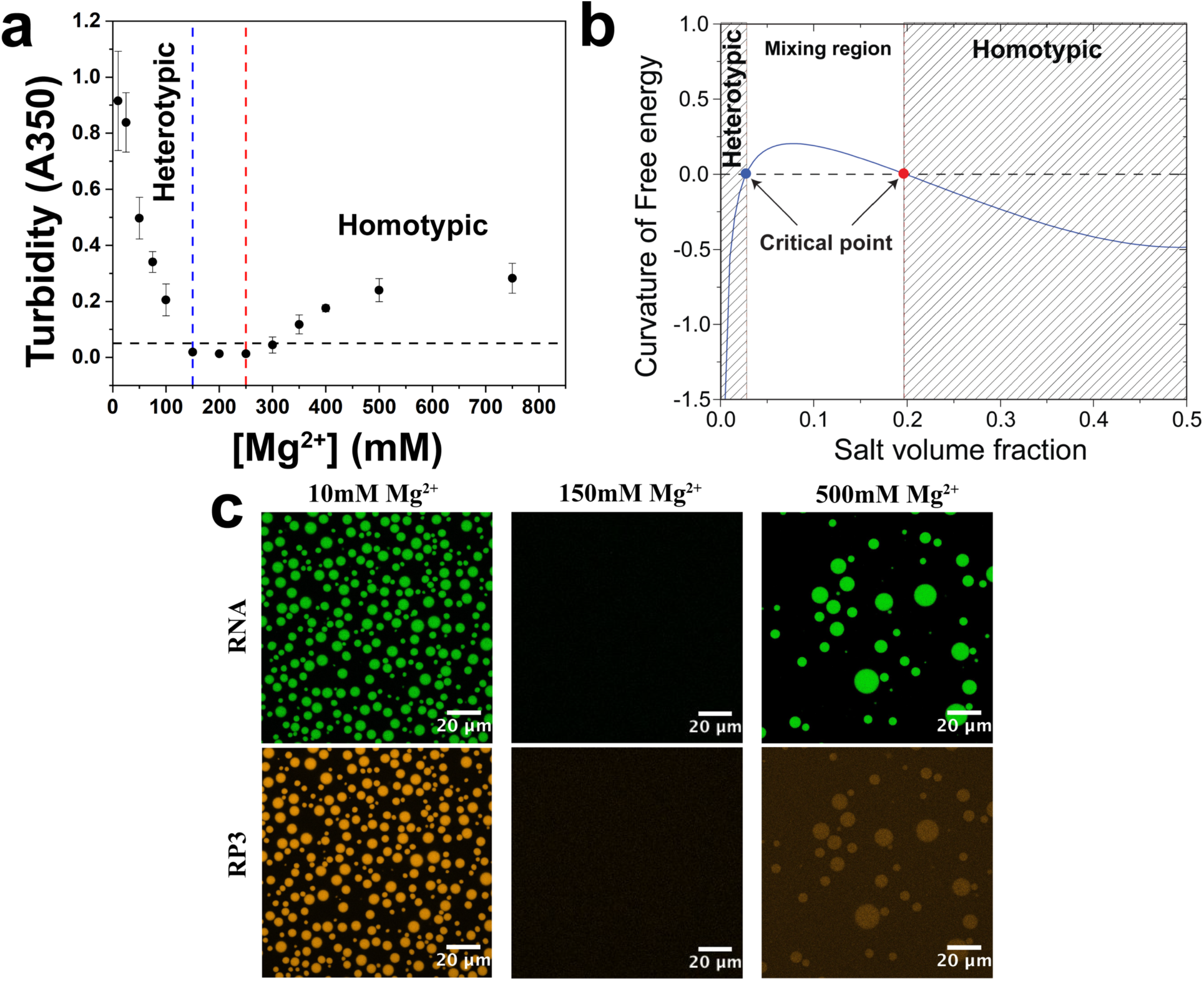
Magnesium drives a switch-like behavior in RNA droplets, creating distinct RNA coacervates. **a.** Solution turbidity measurements of a sequential MgCl_2_ titration showing a window of miscibility (150-250mM MgCl_2_), indicative of droplet switching ([RP3] = 500µM, 2x polyU wt/wt) **b.** Theoretical free energy curvature vs. salt volume fraction calculated using a model (SI Note 1) based on Voorn-Overbeek theory of complex coacervation in the context of Flory-Huggins mean field theory **c.** Confocal fluorescence images of Mg^2+^-dependent droplet switching ([RP3] = 500µM, 2x polyU wt/wt, 0.5µM RP3-AF594, 0.5µM FAM-UGAAGGAC)

To provide a theoretical rationale behind the salt-induced switching behavior of polyU droplet types, we considered a lattice model of thermodynamic free energy. Our free energy expression for the RP_3_-polyU-Mg2+ ternary mixture was derived using Flory-Huggins mean field theory of polymer phase separation and a Debye-Hückel approximation for the ionic interactions^15,38,39^. According to our model, the heterotypic droplets are destabilized by salt since the Debye screening length (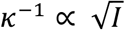; I = ionic strength) decreases with the ionic strength of the solution. In contrast, the homotypic RNA self-assembly is stabilized at higher salt due to (a) a decrease in the repulsive potential of the negatively charged RNA phosphate backbone, and (b) the ability of Mg^2+^ ions to mediate π-stacking of uracil rings^40,41^. Numerical simulation of the free energy curvature, which determines the thermodynamic stability of a mixture with respect to phase separation, indeed showed that the system has two critical points, one at a low salt concentration and one at a higher salt concentration, with a mixed state being populated at intermediate salt concentrations (Figure 2b; SI Note-1). Taken together, our experiments and modeling clearly demonstrate an orthogonal phase behavior of this *in vitro* RNP system in response to a variation in Mg^2+^ concentrations.

### Heterotypic peptide-RNA and homotypic RNA droplets offer distinct microenvironments

Since RNP droplets may function in concentrating cellular proteins and enzymes in an organelle-like microenvironment^42^, any alterations in the selective partitioning of proteins and nucleic acids may be directly linked to their function. Our system requires a more detailed analysis of molecular partitioning given that the fluorescence microscopy images in the droplet switching experiments demonstrate a high preferential inclusion of RP3 within heterotypic RP3-polyU droplets but lower preferential inclusion into homotypic polyU droplets (Fig 2c). Based on a series of confocal imaging experiments, we calculate the estimated partitioning of a range of probes using fluorescent RNA and peptides/proteins as partitioning markers (SI Tables 1-3). We define a partition coefficient as the ratio of fluorescence intensity between the dense and dilute phase (I_dense_/I_dilute_) (Fig S5)^43^. These probes were selected to feature molecular weights ranging from small organic molecules (< 1 kDa) to large globular proteins (~ 80 kDa), along with diversity in charge and structure (Table S4).

Upon increasing cationic concentration, we anticipate an exclusion of RP3 from homotypic polyU droplets due to charge screening of RP3-RNA interactions and a simultaneous strengthening of homotypic polyU interactions (Fig 3a). Experimentally, with increasing Mg^2+^ we observed a continuous decrease in RP3 partitioning into homotypic polyU droplets, eventually leading to preferential exclusion at 1500mM Mg^2+^ (Fig 3a). Although RP3 showed weak partitioning at 500mM Mg^2+^, a peptide (GR)_20_ (20 arginines) with significantly higher positive charge than RP3 (6 arginines) showed favorable partitioning even under these conditions (Fig 3b). However, (GR)_20_ and RP3 partitioning both decreased with increasing Mg^2+^, as expected due to charge screening (Fig 3a, Fig S5, Table S2). Hence, as anticipated based on the overall interaction model, partitioning of positively charged species into homotypic RNA droplets depends on a competition between strengths of peptide-RNA and RNA-RNA interactions, with the higher interaction polyvalency of the (GR)_20_ resulting in a higher degree of competitive peptide-RNA interactions than for RP3. Along with the partitioning decrease, we saw partial selective exclusion of both RP3 and (GR)_20_ at 1500mM Mg_2+_. Additionally, we observed that an archetypal IDP, α-synuclein, positively partitions within heterotypic RP3-polyU droplets (Fig 3b), with electrostatic interactions and its relatively low pI likely being significant factors. However, it is excluded from homotypic polyU droplets (Fig 3b), where the higher ionic strength makes RNA-RNA interactions the dominant competing interaction. Hsp27, a molecular chaperone from the small heat shock protein family also with a pI less than 7, showed similar partitioning properties to that of α-synuclein (Fig 3b), again likely reflecting the same type of competition described above. A large globular fusion protein, eGFP-MBP, with similar pI to the above examples, was observed to be excluded from both types of droplets (Fig 3b). It has been shown that droplet mesh size plays a role in the inclusion/exclusion of biomolecules^44^. One possible explanation to the exclusion of eGFP-MBP is that there is a discrepancy between the size of the protein and the corresponding mesh size of the droplet material. Clear distinctions in partitioning properties between small RNA probes were also observed. U_10_ RNA partitioned well into the RP3-polyU droplets, but poorly into the homotypic polyU droplets (Table S3). A_10_ RNA, however, showed very strong partitioning into polyU droplets. These observed differences can be attributed to the sequence complementarity of A_10_ RNA, which can strongly interact with polyU, as compared to U_10_ RNA. Thus, these differences again reflect a competition in interaction strengths, here however between different types of RNA-RNA interactions. The variation in partitioning among our probes in the homotypic and heterotypic droplets provides additional evidence that these droplets carry biochemically distinct microenvironments.

**Figure 3:**
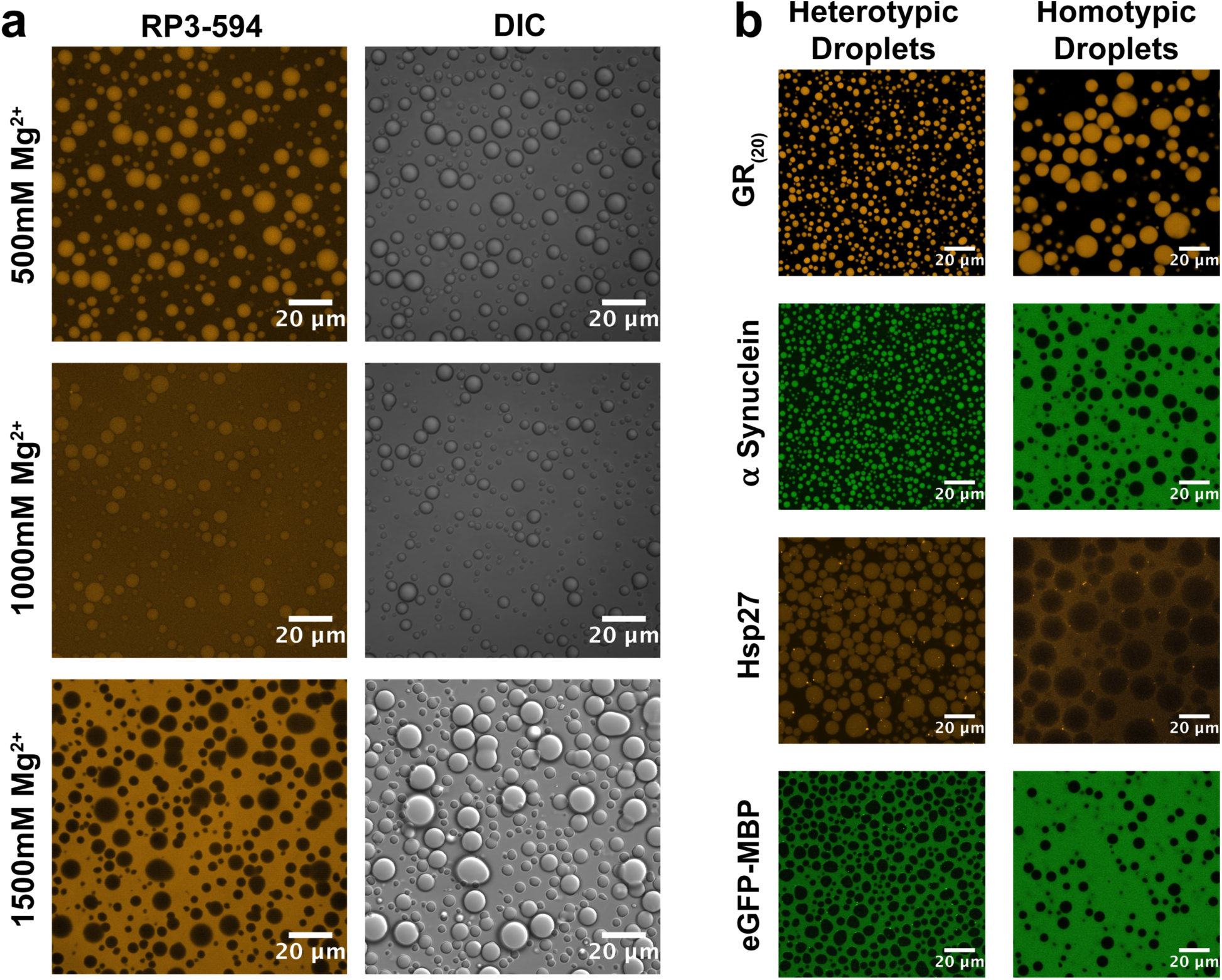
Heterotypic and homotypic droplets display distinct microenvironments. **a.** Confocal fluorescence and DIC images of polyU droplets with RP3-594 as a fluorescent probe (1.5mg/mL polyU, 0.5µM RP3-AF594) **b.** Confocal fluorescence images showing peptide/protein partitioning in heterotypic and homotypic RNA coacervates (Left Panels: [RP3] = 500µM, 0.6x polyU wt/wt, 10mM MgCl_2_ - Right Panels: 1mg/mL polyU, 500mM MgCl_2_) (Fluorescent probes: 0.5µM GR20-AF594, 0.5µM α-synuclein-AF488, 0.5µM Hsp27-AF594, 0.5µM eGFP-MBP-AF488)

### Mg^2+^ controls the material properties of heterotypic and homotypic RNA droplets

The results above show that under certain conditions, a switch-like behavior between heterotypic and homotypic RNA droplets and their differing droplet composition can be controlled by a simple variation in divalent ion concentration. We studied whether the same salt-dependent tuning of the heterotypic and homotypic interactions that are the molecular basis for this switching would also vary the material properties within each type of droplet, though in a more continuous manner. Based on the arguments presented earlier, we would again anticipate opposing effects for the two kinds of droplets upon increasing [Mg^2+^], with a rise in fluidity of heterotypic RP3-polyU droplets and a drop in fluidity in homotypic polyU droplets. To test this idea, we used fluorescence recovery after photobleaching (FRAP) experiments, where the scaled FRAP time (see Figure caption and methods) of an RNA probe (FAM-UGAAGGAC) is taken as a relative measure of molecular diffusivity and hence droplet fluidity under identical experimental conditions.

For RP3-polyU droplets, our results reveal that the recovery time decreases substantially from 0mM Mg^2+^ to 100mM Mg^2+^ (Fig 4a). Additionally, we showed a significant drop in preferential partitioning of this RNA probe between 10mM and 100mM Mg^2+^ (Table S1). The rise in fluidity is consistent in monovalent salt, with an increase in Na^+^ also resulting in a more rapid FRAP recovery (Fig S2). Together, these data show that salt-induced tuning of the electrostatic interactions substantially alters the physical properties of RP3-polyU coacervates. Our observations are broadly consistent with a recent report of salt-induced alteration in protein droplet rheological properties, which follows the same principle^45^.

**Figure 4:**
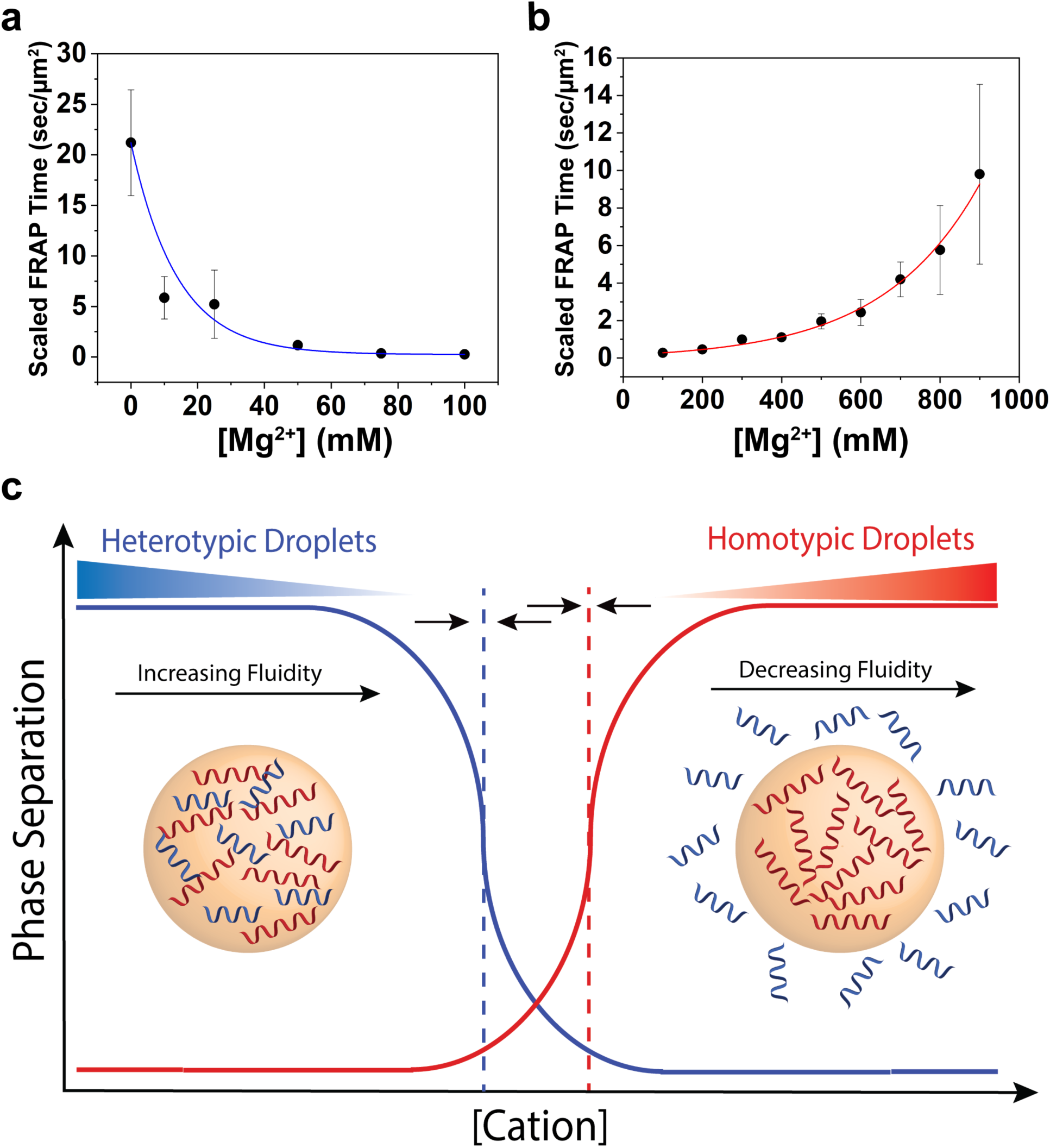
Magnesium concentration alters the fluidity of heterotypic and homotypic droplets. **a.** Scaled FRAP in RP3/polyU droplets as a function of [MgCl_2_]. ([RP3]=500μM, 0.6x polyU wt/wt) The blue line is a visual guide. **b.** Scaled FRAP in polyU droplets as a function of [MgCl_2_] (2 mg/mL polyU, 10% PEG) The red line is a visual guide. **c.** Cartoon representation of switch-like behavior between heterotypic and homotypic droplets as a function of increasing cation concentration. The dotted lines represent the phase boundaries of the distinct droplet types. The arrows indicate that these phase boundaries will move under changing conditions, and are capable of overlapping.

We further probed for the relative fluidity of homotypic polyU droplets as a function of divalent salt concentration. Our predicted trend was observed, in that polyU coacervates undergo progressive hardening with increasing divalent salt concentration (Fig 4b). For example, above a concentration of 900mM Mg^2+^ and in the presence of 10% PEG, the droplet fluidity decreases to a point where the FRAP recovery becomes extremely slow (Fig. S7). This trend is consistent under similar conditions in the absence of PEG (Fig S8). Using the fluorescence intensity of the same RNA probe (FAM-UGAAGGAC), we observed increased RNA partitioning within the droplets as a function of increased [Mg^2+^] (Table S1). These results suggest that at higher concentrations of Mg^2+^ the volume fraction of RNA increases within the droplet phase. Since increasing [Mg^2+^] drives progressive homotypic polyU droplet hardening, we hypothesized that sequestration of Mg^2+^ by a chelating agent would reverse this effect. To test this reversibility, we introduced EDTA in the system, which strongly chelates Mg^2+^ and decreases the bulk concentration of free [Mg^2+^]. Our data revealed that the inclusion of EDTA accelerated the FRAP kinetics (Fig S8). Using microscopy, we show that the addition of EDTA to preformed droplets and subsequent addition of Mg^2+^ can reversibly dissolve and reassemble homotypic polyU droplets (Fig S8). Taken together, these results demonstrate the opposing and reversible effects that Mg^2+^ plays on the fluidity of heterotypic and homotypic droplets.

## Discussion

Homotypic and heterotypic interactions are central in governing the LLPS of RNA-binding proteins containing low-complexity sequences. The feasibility of RNA phase separation by homotypic interactions increases the complexity by which these individual interactions are coupled in regulating the condensed phase dynamics of multi-component RNP granules^46^. In our work, we illuminated some of the aspects of this emerging complexity through the lens of physical and chemical property variations in a simplified model RNP granule. Using experimental results, supported by an analytical model, we showed that a model RNP granule can undergo a switch-like behavior in response to orthogonal RNA interactions. We were able to trigger this behavior in response to a single stimulus, which was the addition of divalent ions. Along with droplet switching, we showed selective and restrictive partitioning of biomolecules, as well as tunable material properties in our distinct condensed phases. Both the switching and tunable behaviors observed in this work can be predicted from theories of phase separation based on the opposing effects of divalent cations on heterotypic and homotypic interactions (Figs 2b, 4c, SI note-1). We anticipate that different conditions could alter the degree of overlap between the two phase-separation regimes, which would instead give rise to continuous tuning behavior. We explored this possibility for the case of Ca^2+^ ions, another important cellular divalent ion. In this case, the lower concentration stabilization of homotypic phase separation results in overlapping regimes and continuous variation in droplet composition (Fig S6, SI note-1).

The single stimulus-driven creation of distinct and tunable condensed phases, with unique chemical and physical properties, reveals potential biological implications. Interaction-based specificity of droplet components can dictate the composition of droplets *in vitro*, suggesting a potential explanation for the extensive variation in composition seen among cellular RNA granules^46–49^. Such a stimulus-dependent modulation of liquid-liquid phase separation may also provide valuable insights into how intracellular RNP granules are regulated in response to versatile environmental cues. As alluded to previously, one motivation for studying divalent cations and their effect on phase separation in our system was the effect of fluctuations in cytoplasmic ion concentrations. Work in the field of ion signaling and homeostasis has shown that changes in ion concentration in cellular compartments are involved in major cellular pathways, such as stress signaling and transcription^20–26^. Variations in cellular Mg^2+^ and Ca^2+^ also have significant relevance in a variety of functions, including timekeeping, neuronal migration and signaling^22–25^. Of special interest to us was the evidence pointing to the rising cytoplasmic concentration of divalent ions during stress, as recent work in phase separation showed that monovalent salt concentration plays a role in stress granule formation^10,50^. Although some of the salt concentrations used in our work might be considered higher than what is found in a cell, crowding may alter this scenario. This is observed in our experiments with PEG-containing buffers, which revealed a substantial reduction in salt requirement. Our work reveals what could be an essential role for divalent ion concentration in this process, tuning not only the material properties of these droplets, but also the accessibility of important biological molecules to specific condensed phases. Specifically, these emergent properties could be necessary in segregation and protection of essential RNAs during cellular stress^51^. We note that the divalent ion-mediated tuning and switching discussed here is potentially one of several mechanisms contributing to the tunability of cellular MLOs. Other types of specific and non-specific interactions between cellular RNA and proteins will therefore be able to alter these effects, providing for a rich tunability and inclusion specificity that can studied in future work. Additionally, understanding the interactions and microenvironments that regulate the phase separation of simple RNAs and peptides could help reveal the necessary driving forces in the formation of protocells, as coacervate biochemistry is a promising direction in deciphering the origin of cells. Lipid membrane vesicles have been proposed as suitable compartments in self-replicating ribozyme systems in a prebiotic RNA world^52^. Homotypic RNA phase separation may provide important advantages as an alternative compartmentalization mechanism. These advantages include stable formation under high divalent ion concentrations needed for ribozyme function^52^, facile exchange of nucleotides, and selective RNA compartmentalization based on specific interactions. Combined, these results reveal a way to consider regulation of LLPS in cellular environments as a complex integration and tuning of weak interactions, which are dependent on but also influential to the cellular microenvironment.

## Materials and Methods

### Peptides/Proteins/RNA Sample Preparation

RP3: ({RRASL}_3_) and RP3C: ({RRASL}_3_C) were ordered and custom synthesized from GenScript (New Jersey, USA) at ≥95% purity. GR20: (C{GR}_20_) was ordered and custom synthesized from PEPSCAN (Lelystad, The Netherlands) at ≥95% purity. RP3 was dissolved in DEPC-treated water (Santa Cruz Biotechnology) and used without further purification. RP3C and GR20 were subsequently fluorescently labeled and purified (See Fluorescent Labeling).

α-synuclein and Hsp27 were expressed and purified in the same manner as our previously published work^53^. *E. coli* (BL21(DE3)) cells were transformed with the eGFP-MBP plasmid. The transformed cells were grown at 37°C to an OD_600_ of ~0.6. Protein expression was induced with IPTG to a final concentration of 0.3mM and allowed to grow for another 4 hours at 37°C. The cultures were harvested by centrifugation and lysed by sonication in lysis buffer (50mM TRIS, 25mM NaCl, 2mM EDTA, pH 8) with a protease inhibitor cocktail. The cell debris was removed by centrifugation and the protein was collected from the supernatant by incubation in Ni-NTA resin. The supernatant was then run through a gravity column, and the protein eluted with an elution buffer containing 250mM imidazole and 150mM NaCl. Dialysis was used to remove imidazole from the buffer. Presence and purity of the protein was checked using SDS-PAGE, A_280_/A_260_ measurements, and mass spectrometry. Polyuridylic acid potassium salt (polyU: 800-1000 kDa) was ordered from Sigma Aldrich (St. Louis, USA). All labeled RNA oligos used as markers in this study ({[6FAM]UGAAGGAC}, {[6FAM]UUUUUUUUUU}, {[6FAM]AAAAAAAAAA}) were ordered from and synthesized by Sigma Aldrich.

### Fluorescent Labeling

*α*-synuclein and Hsp27 were all mono-labeled using cys-maleimide chemistry as described in previous work^53^. α-synuclein was labeled with AlexaFluor488 C5 maleimide (Molecular Probes) and Hsp27 was labeled using AlexaFluor594 C5 maleimide (AF594) (Molecular Probes). The labeling reactions were carried out at 4°C overnight in the dark, and the excess dye was removed by centrifugal filtration with a 3K cutoff filter (Millipore). The purity of the labeled proteins was tested via SDS-PAGE and mass spectrometry. All samples showed a labeling efficiency >90% by UV-Vis measurements.

RP3C and GR20 were labeled in a similar manner with the lyophilized powder resuspended in buffer containing excess AlexaFluor594 C5 maleimide (AF594) (Molecular Probes). The reaction took place overnight at 4°C in the dark. The excess dye was removed by four rounds of acetone precipitation. Four times the sample volume of cold (−20°C) acetone was added to the reaction mixture. The reaction was vortexed and incubated at −20°C for 60 minutes. It was then centrifuged for 10 minutes, and the supernatant disposed of. After four rounds, the acetone was allowed to evaporate and the resulting purified labeled peptide pellet was dissolved in DEPC-treated water (Santa Cruz Biotechnology).

### Turbidity Measurement

All samples were prepared in a buffer background of 10mM TRIS-HCl, pH 7.5. All the turbidity measurements were taken on a NanoDrop 2000c Spectrophotometer (ThermoFisher) at room temperature. For all sequential titration experiments the initial samples (100μL) were prepared to the conditions given in the figure legends. Salt was then added to the solution to the following concentration, the solution was mixed and allowed to equilibrate for 30 seconds, and 3μL were taken out to measure the absorbance at 350nm. For individual points, 5-10μL samples were prepared under diverse salt conditions, mixed, allowed to equilibrate for 30 seconds, and then A_350_ readings were taken. All experimental results were plotted and presented using OriginLab software.

### Microscopy

All DIC and fluorescence microscopy images used in this study were taken on a Zeiss LSM 780 laser scanning confocal microscope. Samples were prepared on Lab-Tek chambered #1.0 borosilicate coverglass. The coverglass was treated with 10% TWEEN 20 for 30 minutes, rinsed with water, and allowed to dry prior to use to avoid sample sticking to the coverslip. Samples were imaged at room temperature using a 63x oil immersion objective (63x oil Plan Apo, 1.4na DIC). All samples, with conditions shown in the figure legends, were prepared with 0.5-1μM of fluorescently labeled markers. For the droplet switching imaging experiment, 2μL of concentrated MgCl_2_ salt solution was injected into the sample during imaging to the given bulk concentration. The same was done for the alternating EDTA/Mg^2+^ experiment. All images and videos were analyzed using Fiji software. Partitioning analysis was done using Zen software. Using the labeled proteins/peptides/RNA as partitioning markers, we made samples under the given conditions with 1μM of labeled samples. Fluorescence intensity was measured within the dense phase and background fluorescence was measured in the dilute phase. The coefficient was calculated as a ratio of droplet intensity over background intensity. At least 15 droplets were analyzed in each sample. In the instances of high partitioning, samples had to be analyzed under increasing laser power to be able to measure background fluorescence. The fluorescence intensity was controlled for saturation, and the partitioning was corrected for laser power. Laser power control experiments were run to ensure that intensity increase was linear with the changes in laser power within the tested range for AF594 and FAM. All dyes (AF488, AF594, FAM) were controlled to ensure that both dye concentrations in the dilute and dense phase were within a linear fluorescence range. Since experimental conditions could result in variations of dye fluorescence quantum yields, which would result in changes to the calculated numbers, we define these numbers as apparent partition coefficients.

### FRAP Experiments

For fluorescence recovery after photobleaching (FRAP) experiments, the samples were prepared in the same manner as described above with conditions given in the figure legends. The {[6FAM]UGAAGGAC} RNA oligo was used as a marker for all FRAP experiments. For all FRAP images, one droplet was bleached (with a ROI between 3-25 µm^2^) and one was used as a reference marker. Experimental droplets were bleached using 10 iterative pulses of 100% laser power. Both the fluorescence recovery of the bleached droplet, and the changing fluorescence of the reference droplet due to photobleaching were collected and analyzed. The reference droplet was used as a baseline to calculate a normalized intensity. The normalized curves were plotted and fitted on OriginLab Software using an exponential fit (*y* = *y*_0_ + *A*_1_*e*^*x*/*τ*^), from which the τ value (time constant) was obtained. Finally, the τ value was normalized based on the bleached ROI of the droplets (τ/ROI) to give the scaled FRAP time. Multiple droplets (n = 3-6) were used at each salt condition to extract a mean scaled FRAP time which is presented with error bars representative of one standard deviation.

## Acknowledgements

We gratefully acknowledge support for this work from NIGMS/NIH (Grants RO1 GM066833, RO1 GM115634 and R35 GM130375 to A.A.D.) and National Science Foundation (Grant MCB 1818385 to A.A.D.) and from University at Buffalo, SUNY, College of Arts and Sciences to P.R.B. We thank Ian J. Macrae, Irem Nasir, Emily Bentley, and Jose Luis Olmos Jr. for valuable feedback regarding this work. We thank Emily Bentley for providing the eGFP-MBP used in the study.

## Author contributions

P.L.O., A.N.M., P.R.B, and A.A.D. designed the study. P.L.O., A.N.M., and P.R.B. designed the experimental strategies. P.L.O. and A.N.M. collected and analyzed all turbidity, confocal microscopy, partitioning, and FRAP data. P.R.B. generated the recombinant proteins and performed protein fluorescence labeling. P.L.O. and A.N.M. performed peptide fluorescence labeling. I.A. constructed the analytical model and performed the free energy surface simulation. All authors contributed in writing the manuscript.

