## Supplementary Information for "Divalent cations can control a switch-like behavior in heterotypic and homotypic RNA coacervates"

**Supplementary Materials**

**SI-note 1**

To model the phase behavior of RP3-poly(U)-$Mg^{2+}$ mixture, we consider the Voorn-Overbeek theory of complex coacervation in the context of Flory-Huggins mean field theory of polymer phase separation[1,2]. For practical purposes, we make a simplified assumption that our system is composed of a polyelectrolyte salt, PR, and an ionic salt, $MgCl_2$. In our model, P is a polycation and Q is a polyanion, having the same size N and the same charge density $\sigma$. To ensure electro-neutrality, the two polymers must have equal volume fraction ($\phi_P = \phi_R$). The key idea of this model is to map the complex coacervation, which is an *associative phase separation*, into a *segregative phase separation*. As a starting point, we consider the Voorn-Overbeek free energy of mixing per lattice site, which is given by:

$$\frac{F}{kT} = \frac{\phi}{N}\ln\frac{\phi}{2} + c\ln c + (1-\phi-c)\ln(1-\phi-c) - \alpha(\sigma\phi + c)^{\frac{3}{2}} \qquad (1)$$

Where $\phi = \phi_P + \phi_R$ is the volume fraction of the polyelectrolyte salt and c is the ionic salt volume fraction. The effect of salt is considered here by an additional term in the translational entropy as well as the enthalpy part of equation-1. This equation can be mapped into an effective Flory interaction parameter of a polymer-solvent system that phase separates by *segregative mechanism* as described by: [3]

$$\chi_{eff} = \chi + \chi_{ion} = \chi + \frac{2\pi}{3}\frac{l_B^2\kappa^{-1}\sigma^2}{l^3} \qquad (2)$$

Where $\kappa^{-1}$ is the Debye length, $l_B$ is the Bjerrum length and $l$ is the size of the monomeric unit in the polyelectrolyte. The first term represents the short-range excluded volume interaction, which is set to zero ($\chi$=0) for simplicity, as in the case of an athermal solvent[4]. Consequently, the corresponding free energy density equation in the Flory-Huggins framework is given by:

$$\frac{F}{kT} = \frac{\phi}{N}\ln\frac{\phi}{2} + (1-\phi-c)\ln(1-\phi-c) + c\ln c + \chi_{eff}\phi(1-\phi-c) \qquad (3)$$

In equation-3, the entropic contribution of salt can be considerable since the salt concentrations used in the experiment are relatively high. The Debye length can be expressed in terms of the ionic strength *I* as described in: [5]

$$\kappa^{-1} = \sqrt{\frac{\epsilon_r\epsilon_0 kT}{2x10^3 N_A e^2 I}} \qquad (4)$$

Where I is proportional to the number concentration of ions $\psi$ as $I = \frac{1}{2}\sum_{i=1}^{n}\psi_i z_i^2 \propto \psi = \frac{c}{\nu}$. $\nu$ is the volume of a single ion. For $MgCl_2$; $I = \frac{5}{2}\psi = \frac{5}{2}\frac{c}{\nu}$. Rewriting equation-2 in terms of salt volume fraction, we get

$$\chi_{eff} = \chi_{ion} = \frac{\beta}{\sqrt{c}} \qquad (5)$$

Where $\beta = \frac{2\pi}{3}\frac{l_B^2\sigma^2}{l^3}\sqrt{\frac{2\epsilon_r\epsilon_0 kT\nu}{10^4 N_A e^2}}$. Equation-3 now becomes,

$$\frac{F}{kT} = \frac{\phi}{N}\ln\frac{\phi}{2} + (1-\phi-c)\ln(1-\phi-c) + c\ln c + \frac{\beta}{\sqrt{c}}\phi(1-\phi-c) \qquad (6)$$

Equation-6 is sufficient to describe the phase behavior of a system that phase separates at low salt concentration and but undergoes mixing at higher salt concentration. To account for homotypic condensation of the polyanion, R (*i.e.,* RNA in our case), at higher salt concentration, we consider a term for the enthalpy coming from *salt-R* interactions that satisfy two conditions: (i) it must be proportional to the square of the volume fraction of R molecules $\left(\frac{\phi}{2}\right)^2$ in order to contribute to the curvature of the free energy $\frac{\partial^2 F}{\partial\phi^2}$, and (ii) it should be a function of salt concentration since our FRAP data indicate increased attraction between RNA molecules with increasing [$MgCl_2$]. These conditions lead us to consider a three body interaction term: [6]

$$\delta F_{3-body} = -\frac{\epsilon c\phi^2}{4} \qquad (7)$$

Where $\epsilon$ is the strength of the interaction. This interaction may emerge from divalent cations mediating $\pi$-stacking of uracil rings in the case of poly(U) RNA [7,8]. Finally, the total free energy can be written as:

$$\frac{F}{kT} = \frac{\phi}{N}\ln\frac{\phi}{2} + c\ln c + (1-\phi-c)\ln(1-\phi-c) + \frac{\beta}{\sqrt{c}}\phi(1-\phi-c) - \frac{\epsilon c\phi^2}{4} \qquad (8)$$

The stability of the mixed state is determined by the curvature of the free energy, a positive curvature indicates a locally stable homogeneous phase, while a negative curvature indicates phase separation[9]. A spinodal boundary of the system is constructed by calculating the inflection points of the free energy equation ($\frac{\partial^2 F}{\partial\phi^2} = 0$) and shown in Fig S1a. At low salt volume fraction, the free energy curvature is negative and the homogeneous phase is thermodynamically unstable. Therefore, the system spontaneously separates into *PR-rich* and *PR-poor* phases. Increasing the salt concentration causes the two-phase region to disappear and the curvature becomes positive, hence a homogeneous phase emerges. Interestingly, the curvature becomes negative again upon further increase in salt concentration, leading the system to phase separate due to contributions from the three body interaction term (Fig S1b). This is consistent with the experimentally obtained phase boundary curves of RP3-poly(U)-$Mg^{2+}$ mixture, which is shown in Fig 2a.

In conclusion, here we presented a simple phase separation model which qualitatively captures the essential features of our ternary mixture phase behavior. Our model also predicts that for higher values of the parameter $\beta$, which is proportional to the square of linear charge density, the mixing region should vanish and the free energy curvature should remain negative throughout the range of salt concentrations (Fig S1b). This scenario can be realized for polycations containing significantly higher number of arginine residues than our present peptide system.

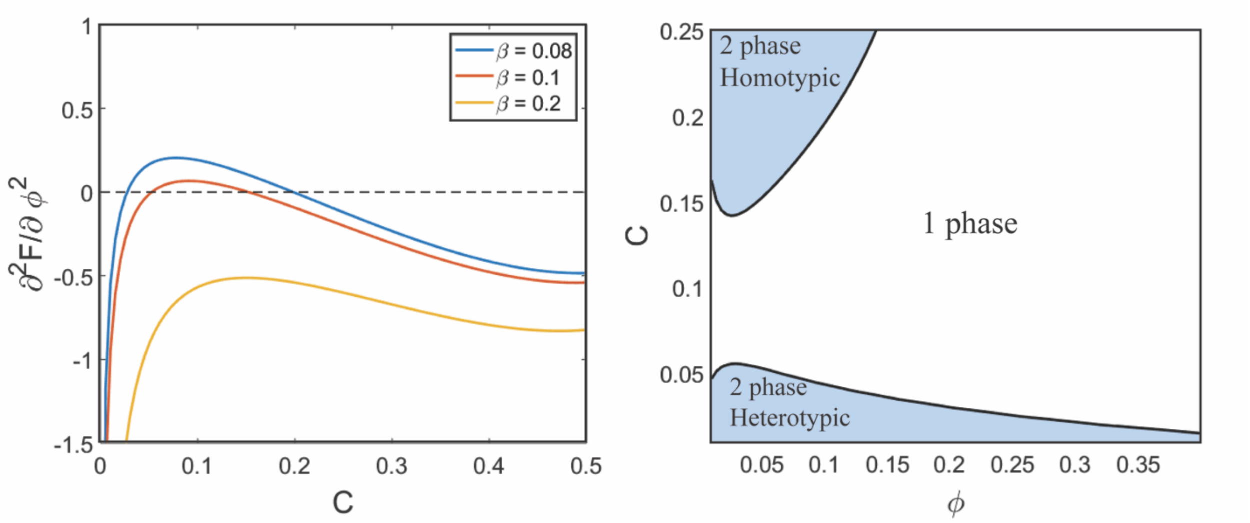


**Figure S1: a.** Spinodal boundary curve calculated using equation-8. Parameters used: N=1000, $\beta$ = 0.1 and $\epsilon = 10$ **b.** Free energy curvature $\frac{\partial^2 F}{\partial \phi^2}$ as a function of salt concentration for the same parameters at volume fraction $\phi = 0.05$ and variable $\beta$, which is directly related to square of linear charge density.

**Supplementary Figure 2**

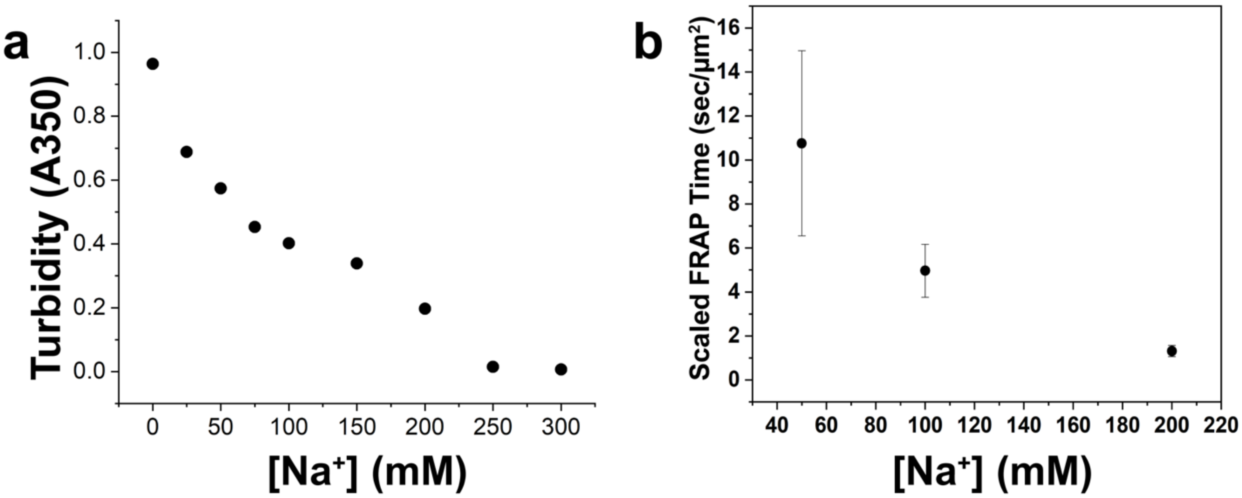


**Figure S2: Sodium concentration leads to decreased formation and variation of RP3-polyU droplet fluidity. a.** Solution turbidity measurements of RP3-polyU droplets as a function of [NaCl]. ([RP3] = 500µM, 0.6x polyU wt/wt) **b.** Scaled FRAP in RP3-polyU droplets as a function of [NaCl] ([RP3] = 500µM, 0.6x polyU wt/wt).

**Supplementary Figure 3**

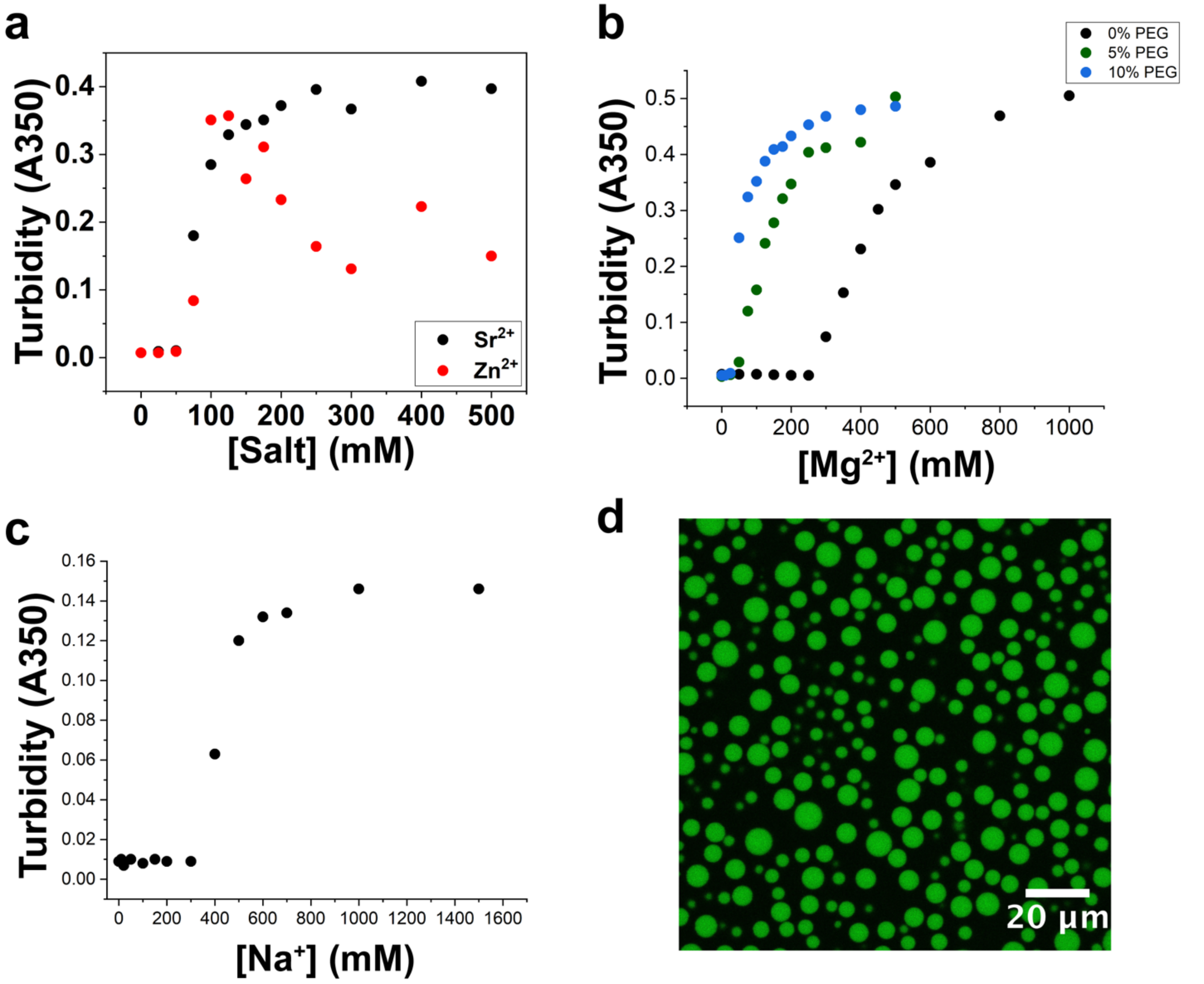


**Figure S3: PEG influences the phase boundaries in the presence of divalent and monovalent salts. a.** Solution turbidity measurements of polyU droplets as a function of [$SrCl_2$] and [$ZnCl_2$] (1mg/mL polyU) **b.** Solution turbidity measurements of polyU droplets as a function of [$MgCl_2$] in the presence of PEG8000 (1mg/mL polyU, PEG8000) **c.** Solution turbidity measurements as a function of [NaCl] (1.5 mg/mL polyU, 10% PEG) **d.** Confocal fluorescence image showing polyU droplets in the presence of NaCl (2mg/mL polyU, 1M NaCl, 10% PEG8000).

**Supplementary Figure 4**

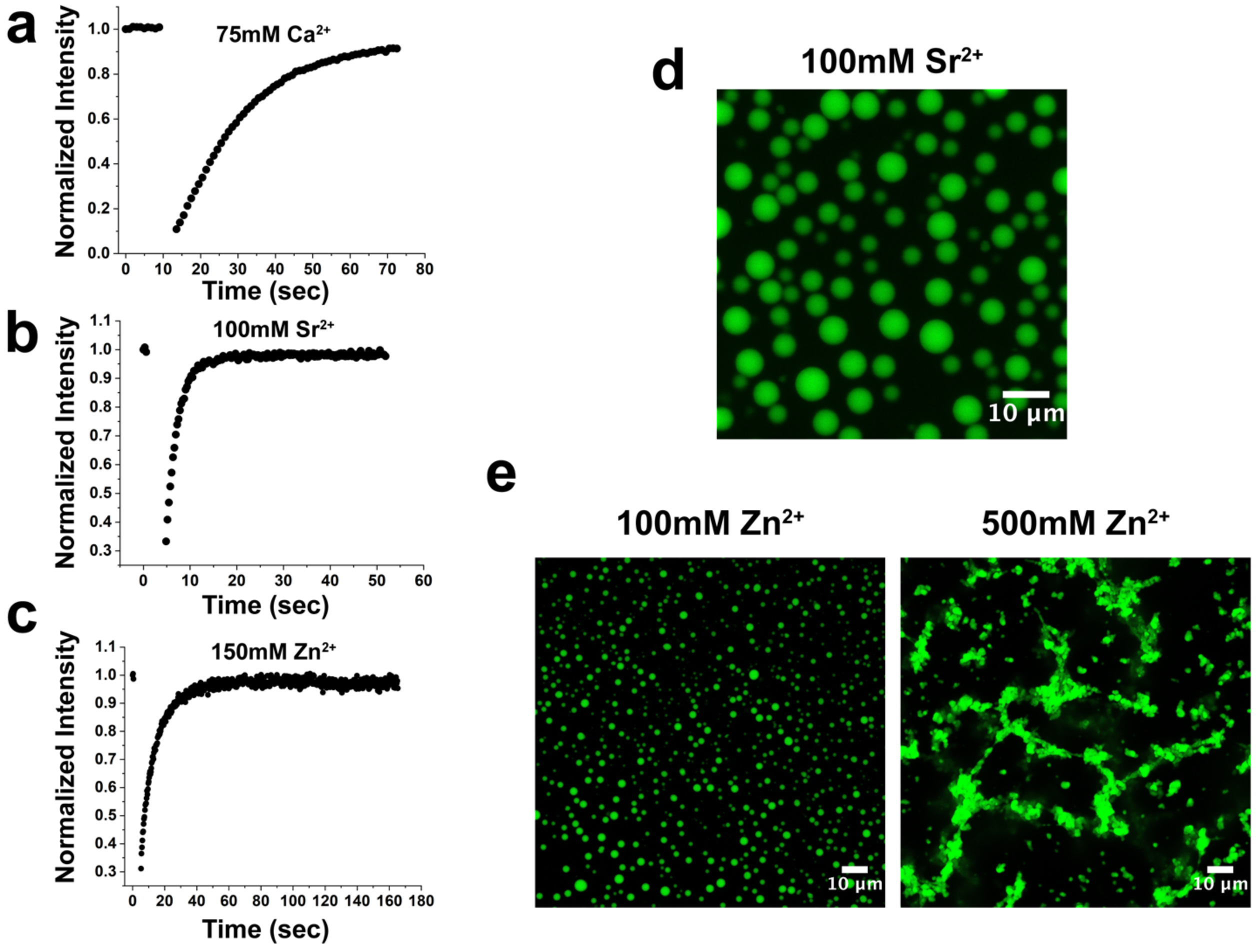


**Figure S4: Multiple divalent cations drive formation of fluid homotypic polyU droplets a.** FRAP curve of homotypic polyU droplets showing fluidity and full recovery (75mM $CaCl_2$, 2mg/mL polyU) **b.** FRAP curve of homotypic polyU droplets showing fluidity and full recovery (100mM $SrCl_2$, 1mg/mL polyU) **c.** FRAP curve of homotypic polyU droplets showing fluidity and full recovery (150mM $ZnCl_2$, 1mg/mL polyU) **d.** Confocal fluorescence image showing polyU droplets in the presence of $SrCl_2$ (1mg/mL polyU, 100mM $SrCl_2$) **e.** Confocal fluorescence images showing polyU droplets in the presence of 100 mM $ZnCl_2$ (left panel) and polyU aggregates in the presence of 500mM $ZnCl_2$ (right panel)(1mg/mL polyU in both panels).

**Supplementary Figure 5**

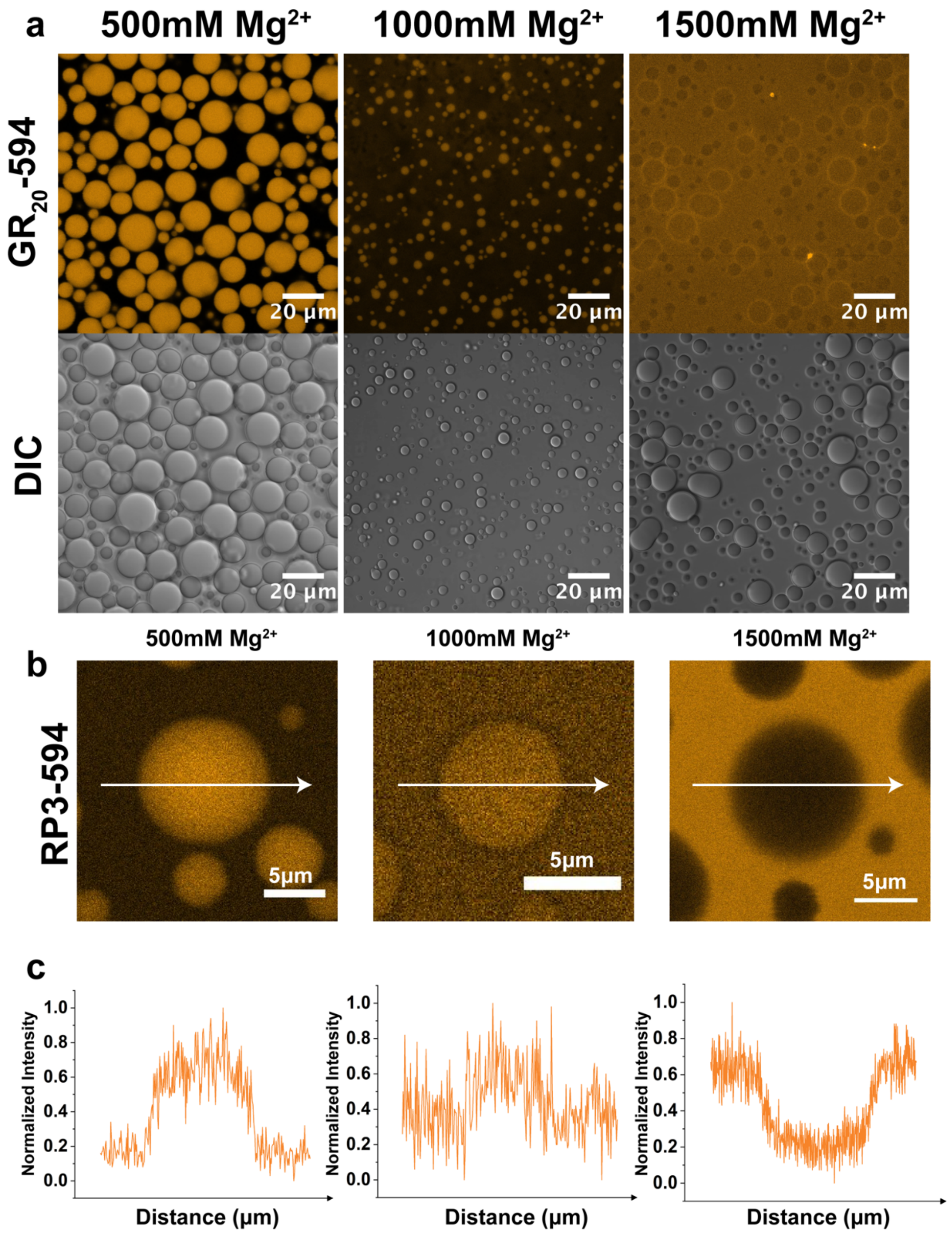


**Figure S5: PolyU droplets exclude positively-charged peptides at high $Mg^{2+}$ concentration.** **a.** Confocal fluorescence and DIC images of polyU droplets with Alexa594-labeled GR20 as a fluorescent probe. ([polyU] = 1.5mg/mL, 0.5µM GR20-AF594). **b.** Confocal fluorescence images showing Alexa594-labeled RP3 partitioning into homotypic polyU droplets ([polyU] = 1.5mg/mL, 0.5µM RP3-AF594). **c.** Fluorescence intensity profiles of the images in b., with the distance axis corresponding to the white arrows.

**Supplementary Figure 6**

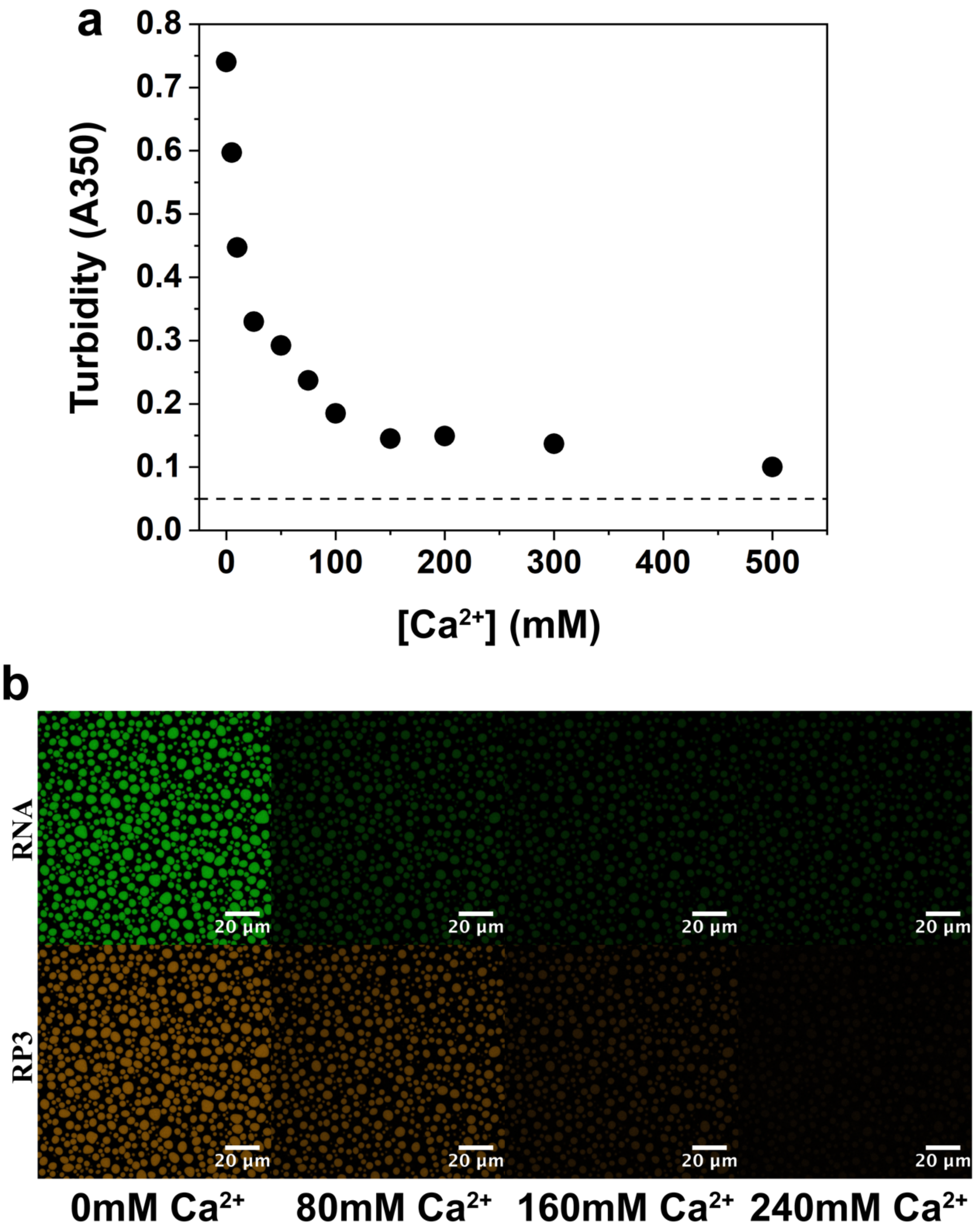


**Figure S6: Heterotypic and homotypic droplets can overlap, showing no window of miscibility a.** Solution turbidity measurements of RP3-polyU droplets as a function of [$CaCl_2$]. The dotted line represents our established turbidity cutoff for phase separation at $A_{350}$ = 0.05 ([RP3] = 500µM, 0.6x polyU wt/wt). **b.** Confocal fluorescence microscopy images of droplets in a sequential titration of $CaCl_2$. Predicted concentration of $Ca^{2+}$ is shown below the frames. ([RP3] = 500µM, 0.6x polyU wt/wt, 0.5µM FAM-UGAAGGAC, 0.5µM RP3-AF594).

**Supplementary Figure 7**

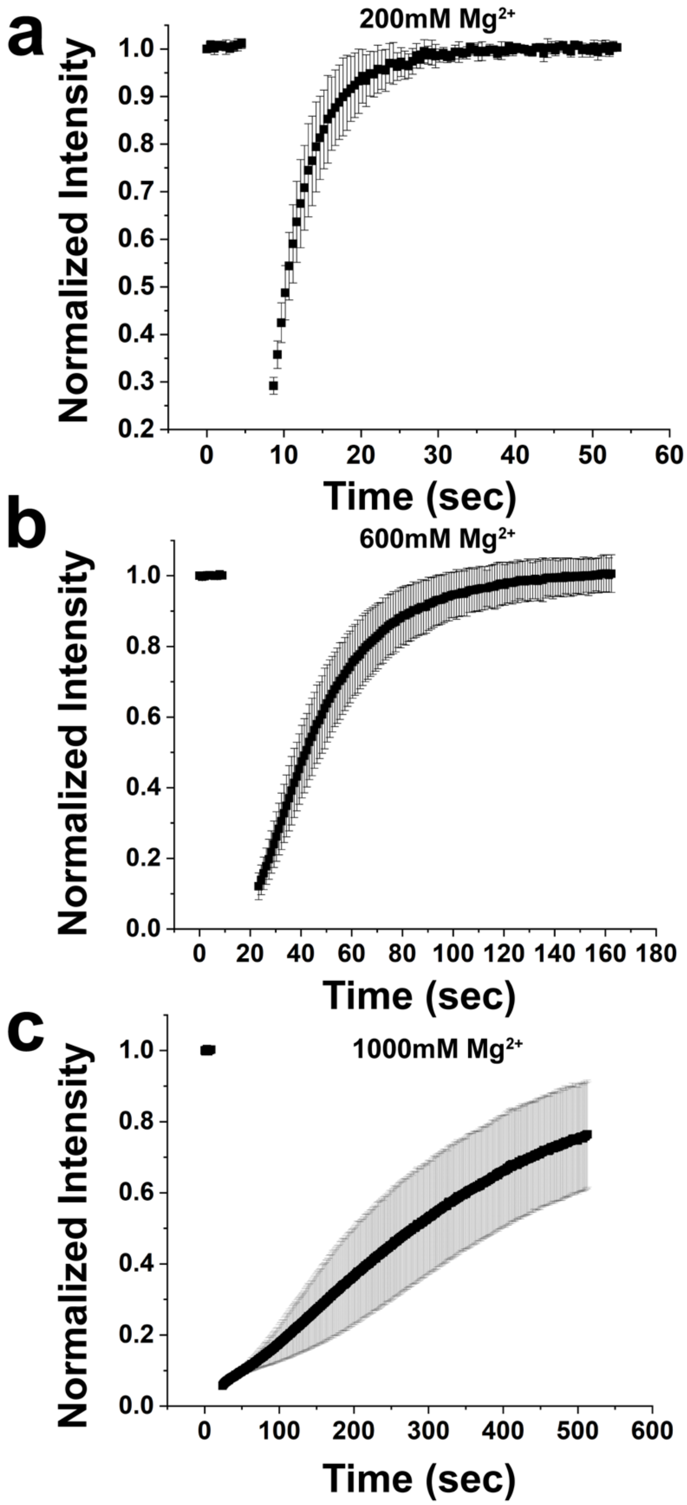


**Figure S7: Fluorescence recovery after photobleaching at varying $Mg^{2+}$ concentrations. a.** 200mM $Mg^{2+}$ ([polyU] = 2 mg/ml, 10% PEG) **b.** 600mM $Mg^{2+}$ ([polyU] = 2 mg/ml, 10% PEG) **c.** 1000mM $Mg^{2+}$ ([polyU] = 2 mg/ml, 10% PEG).

**Supplementary Figure 8**

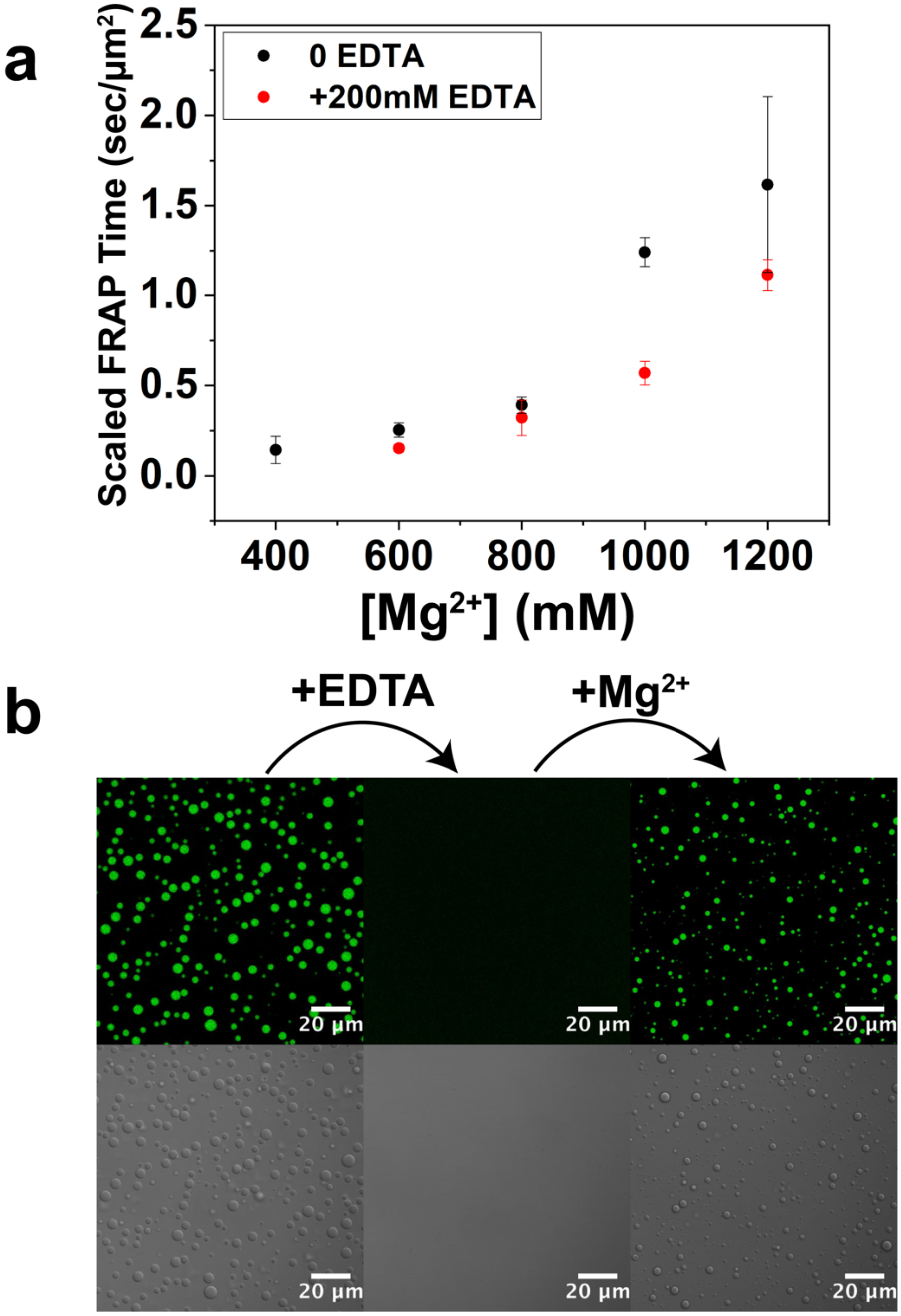


**Figure S8: EDTA reverses the effect of $Mg^{2+}$ on homotypic polyU droplets a.** Scaled FRAP times in the absence and presence of EDTA (2mg/mL polyU) **b.** Fluorescence and DIC microscopy images showing reversible dissolution and formation of polyU droplets by consecutive addition of EDTA and $MgCl_2$ (Frame1: 1.5mg/mL polyU, 35mM $MgCl_2$, 10% PEG, 0.5µM FAM-UGAAGGAC - Frame 2: Addition of 50mM EDTA - Frame 3: Addition of 100mM $MgCl_2$).

### Estimated Partitioning Coefficient

**Table 1**

| | 10mM $Mg^{2+}$, 500µM RP3, .6X polyU | 100mM $Mg^{2+}$, 500µM RP3, .6X polyU | 400mM $Mg^{2+}$, 1mg/mL polyU | 1000mM $Mg^{2+}$, 1mg/mL polyU |
|---|---|---|---|---|
| [6FAM]UGAAGGAC | **2438 ± 422** | **142 ± 10** | **57 ± 3.7** | **91 ± 4.3** |

**Table 2**

| | 10mM $Mg^{2+}$, 500µM RP3, .6X polyU | 500mM $Mg^{2+}$, 1mg/mL polyU | 1000mM $Mg^{2+}$, 1mg/mL polyU | 1500mM $Mg^{2+}$, 1mg/mL polyU |
|---|---|---|---|---|
| RP3-AF594 | **11386 ± 1887** | **2.9 ± 0.1** | **1.4 ± 0.04** | **0.77 ± 0.03** |
| GR20-AF594 | **5721 ± 767** | **65 ± 13** | **3.8 ± 0.46** | **0.95 ± 0.03** |

**Table 3**

| | 10mM $Mg^{2+}$, 500µM RP3, .6X polyU | 500mM $Mg^{2+}$, 1mg/mL polyU |
|---|---|---|
| [6FAM]U10 | **759 ± 144** | **2.3 ± 0.1** |
| [6FAM]A10 | **N/A** | **1134 ± 95** |
| AF488 | **39 ± 2.3** | **1.3 ± 0.04** |
| EGFP-MBP-AF488 | **0.39 ± 0.015** | **0.095 ± 0.0088** |
| α-synuclein-AF488 | **4.4 ± 0.17** | **0.13 ± 0.0057** |
| HSP27-AF594 | **14 ± 2.1** | **0.72 ± 0.083** |

These tables represent the apparent partitioning estimated from fluorescence intensity measurements of droplets and the background, as described in the Microscopy section of the Materials and Methods. This same procedure is widely used in the field. The error (1 standard deviation) was determined based on several intensity measurements for each sample. While additional considerations can apply to obtaining absolute partition coefficients (see Materials and Methods), the numbers presented here provide a reliable estimate within the order of magnitude for preferential partitioning vs. exclusion, which were used for our conclusions in the paper.

**Table 4 – Proteins/Peptides Molecular Weight and Theoretical PI**

| | **Molecular Weight (kDa)** | **Theoretical pI** |
|---|---|---|
| EGFP-MBP | 77.6 | 5.03 |
| α-synuclein | 14.5 | 4.67 |
| HSP27 | 22.8 | 5.98 |
| RP3 | 1.9 | 12.3 |
| GR20 | 4.4 | 12.95 |
